# Eliminating effects of particle adsorption to the air/water interface in single-particle cryo-electron microscopy: Bacterial RNA polymerase and CHAPSO

**DOI:** 10.1101/457267

**Authors:** James Chen, Alex J. Noble, Jin Young Kang, Seth A. Darst

## Abstract

Preferred particle orientation presents a major challenge for many single particle cryoelectron microscopy (cryo-EM) samples. Orientation bias limits the angular information used to generate three-dimensional maps and thus affects the reliability and interpretability of the structural models. The primary cause of preferred orientation is presumed to be due to adsorption of the particles at the air/water interface during cryo-EM grid preparation. To ameliorate this problem, detergents are often added to cryo-EM samples to alter the properties of the air/water interface. We have found that many bacterial transcription complexes suffer severe orientation bias when examined by cryo-EM. The addition of non-ionic detergents, such as NP-40, does not remove the orientation bias but the Zwitter-ionic detergent CHAPSO significantly broadens the particle orientation distributions, yielding isotropically uniform maps. We used cryoelectron tomography to examine the particle distribution within the ice layer of cryo-EM grid preparations of *Escherichia coli* 6S RNA/RNA polymerase holoenzyme particles. In the absence of CHAPSO, essentially all of the particles are located at the ice surfaces. CHAPSO at the critical micelle concentration eliminates particle absorption at the air/water interface and allows particles to randomly orient in the vitreous ice layer. We find that CHAPSO eliminates orientation bias for a wide range of bacterial transcription complexes containing *E. coli* or *Mycobacterium tuberculosis* RNA polymerases. Findings of this study confirm the presumed basis for how detergents can help remove orientation bias in cryo-EM samples and establishes CHAPSO as a useful tool to facilitate cryo-EM studies of baterial transcription complexes.

## Introduction

Advances in single particle cryo-electron microscopy (cryo-EM) now allow structure determination of biological macromolecular complexes to near atomic-resolution [1]. Nevertheless, a major complication for many biological cryo-EM specimens is particle orientation bias [2,3]. Specimens that suffer particle orientation bias can have an anisotropic distribution of angular projection directions leading to under-sampling of Fourier components in the final three-dimensional reconstruction. This under-sampling leads to an overall loss of structural information parallel to the axis of preferred orientation, giving the maps an anisotropic point spread function leading to a “smearing effect” artifact [4-6], which affects the interpretability of the cryo-EM maps.

In examining a complex of *Escherichia coli* (*Eco*) 6S RNA bound to RNAP σ^70^-holoenzyme (6S-Eσ^70^) for single-particle cryo-EM structure determination, we encountered a severe orientation bias problem [7]. We explored a range of solution conditions and detergent additives to solve the orientation bias problem and discovered that the detergent 3-([3-Cholamidopropyl]dimethylammonio)-2-hydroxy-1propanesulfonate (CHAPSO) was uniquely effective. We hypothesized that the preferred orientation was due to adsorption and orientation of the particles at the air-water interface [8] and that CHAPSO mitigated this problem by preventing adsorption at the interface. We used fiducial-less cryo-electron tomography (cryo-ET) on the single-particle specimens to visualize particle distributions within the vitreous ice [9,10]. The results confirmed our hypothesis; the orientation bias arises from interactions of the particles with the surfaces of the ice layer. In the presence of a sufficient concentration of CHAPSO, the particles were excluded from the ice surfaces and distributed within the ice layer with nearly random orientations. We show that CHAPSO solves preferred orientation problems for a number of single-particle samples comprising bacterial transcription complexes.

## Results

### 6S-Es^70^ particles show severe orientation bias which is significantly relieved with CHAPSO

Initial single particle cryo-EM analysis of the 6S-Eσ^70^ complex was performed using a potassium L-glutamate (KGlu, Supplement Table 1). Micrographs showed uniform, homogenously dispersed particles that yielded detailed high-resolution 2D classes (Supplement Fig. 1). However, 3D alignment resulted in “smeared” maps, suggesting that the particles in the vitreous ice layer exhibited orientation bias (Supplement Fig. 1). The angular distribution plot of the particles aligned to a low-pass filtered X-ray crystal structure of *Eco* core RNAP (PDB ID 4LJZ with σ^70^ removed; [11]) as a template revealed a distribution corresponding to essentially one orientation (Fig. 1A). A rough characterization of the particle distribution as a Gaussian yielded a peak at about rotation angle (rot) of −19°, tilt angle (tilt) −13°, and a standard deviation (±) of 15°.

**Fig. 1.**
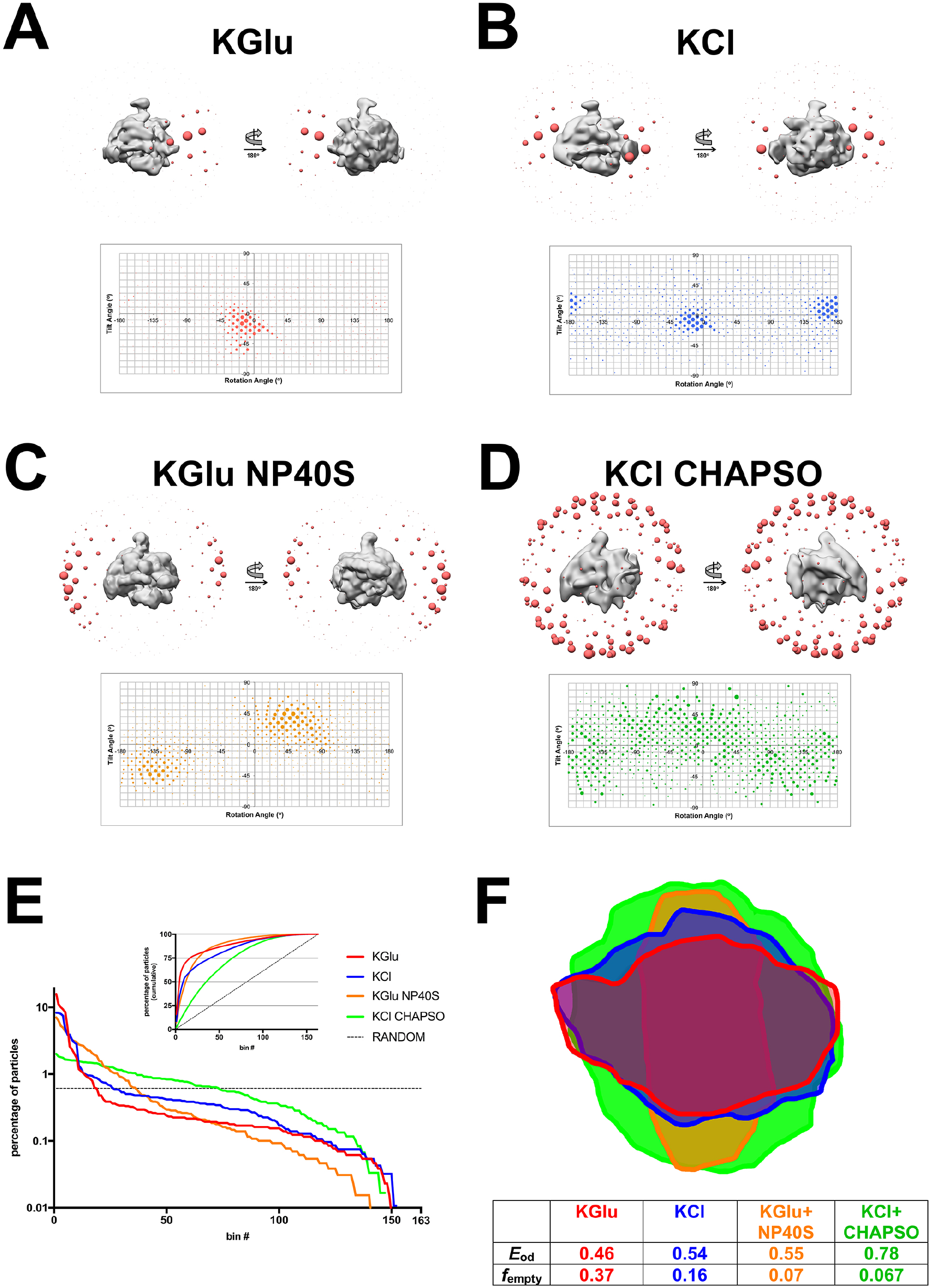
Single particle cryo-EM analysis of 6S-Eσ^70^ particle orientation distributions in different solution conditions. A – D. (Top Panel) 3D distribution plot of particle orientations. Particles were 3D classified into one class using *Eco* core RNAP (PDB ID 4LJZ [14], σ^70^ was deleted and the structure was low-pass filtered using EMAN2) [35] as a 3D template in RELION-2.1 [34]. The resulting density is shown as a solid grey volume and the angular distribution from this alignment is shown as red spheres. Each sphere represents a particular Euler angle and the sphere volume represents the absolute number of particles at that particular angle. (Bottom Panel) 2D distribution plot of particle orientations. Particles are plotted on a tilt angle vs rotation angle graph. Areas of the points represents the percentage of particles at that particular orientation. A. Angular distribution of 6S-Eσ^70^ particles in KGlu (see Supplementary Table 1). B. Angular distribution of 6S-Eσ^70^ particles in KCl. C. Angular distribution of 6S-Eσ^70^ particles in KGlu+NP40S. D. Angular distribution of 6S-Eσ^70^ particles in KCl+CHAPSO. E. Particles for each dataset were grouped into Euler angle bins (20° rotation angle x 20° tilt angle bind) and then the bins were ranked according to the number of particles populating that bin (bin #1 has the most particles, so on). Plotted on a semi-log scale is the percent of total particles in each dataset by bin #. The horizontal dashed line represents a totally random particle orientation distribution (equal number of particles in each bin). (Inset) Plotted is the cumulative percent particles by bin #. The random distribution is denoted by the dashed line. F. Cross-sections through the middle of the expected PSFs (calculated using cryoEF [5]) are superimposed, illustrating the anisotropy for the KGlu (red), KCl (blue), and KGlu+NP40S (orange) samples, while the KCl+CHAPSO sample yields an isotropic PSF (green). Parameters further characterizing the orientation distributions (the orientation efficiency, *E*_od_, and the fraction of unsampled Fourier space, *f*_empty_ [5]) are also tabulated.

To overcome this particle orientation bias, we prepared and analyzed samples in a number of alternative conditions, such as a buffer containing KCl instead of KGlu (KCl, Supplemental Table 1), and the original KGlu condition but with the addition of the non-ionic detergent Nonidet P40 substitute (KGlu-NP40S, Supplemental Table 1).

Single particle cryo-EM analysis of the 6S-Eσ^70^ particles in the KCl condition also showed severe orientation bias (Fig. 1B), with one peak at an orientation and spread very similar to the KGlu condition (rot −12°, tilt −8°, ± 14°), but with an additional peak at approximately (rot 168°, tilt 8°), corresponding to a mirror image projection of the first orientation. Since the mirror image projection does not contribute any new information to the 3D reconstruction, having two mirror image orientations is equivalent to having one orientation. Preparation of the particles in KGlu+NP40S also yielded two mirror image peaks [(rot −130°, tilt −47°) and (rot 50°, tilt 47°)]. The distribution was broadened with respect to the KGlu and KCl distributions, with standard deviation ± 20°. Thus, the addition of the detergent still gave only one effective orientation, but the bias was slightly ameliorated. In contrast to KGlu, KCl, and KGlu+NP40S, particles prepared in KCl+CHAPSO did not exhibit peaks of preferred orientation; instead the particle orientations were spread over a large fraction of Euler angles (Fig. 1D), resulting in isotropically uniform 3D reconstructions.

For each individual sample, the particles were grouped into 163 bins according to their Euler angles (corresponding to 20° increments in rotation and tilt angles) and ranked according to the percentage of particles in each bin (bin #1, highest %; bin #2, next highest %; so on). The orientation distribution of the particles prepared in each condition was compared by plotting the histograms of the % particles in each bin according to bin # (Fig. 1E). A completely random distribution of particle orientations would yield a flat distribution, with 0.61% particles in each bin (dashed horizontal line in Fig. 1E). Visualizing the orientation distributions this way highlights the bias of the KGlu, KCl, and KGlu+NP40S samples. The KCl+CHAPSO sample, while not completely randomized, approaches the random orientation distribution more closely. The inset of Fig. 2E plots the cumulative % of particles across the bins. This plot reveals that 50% of the particles are binned into only 5, 9 and 12 bins for the KGlu, KCl, and KGlu+NP40S conditions, respectively, while 50% of the KCl+CHAPSO particles are spread out over 36 bins (Fig. 1E).

**Fig. 2.**
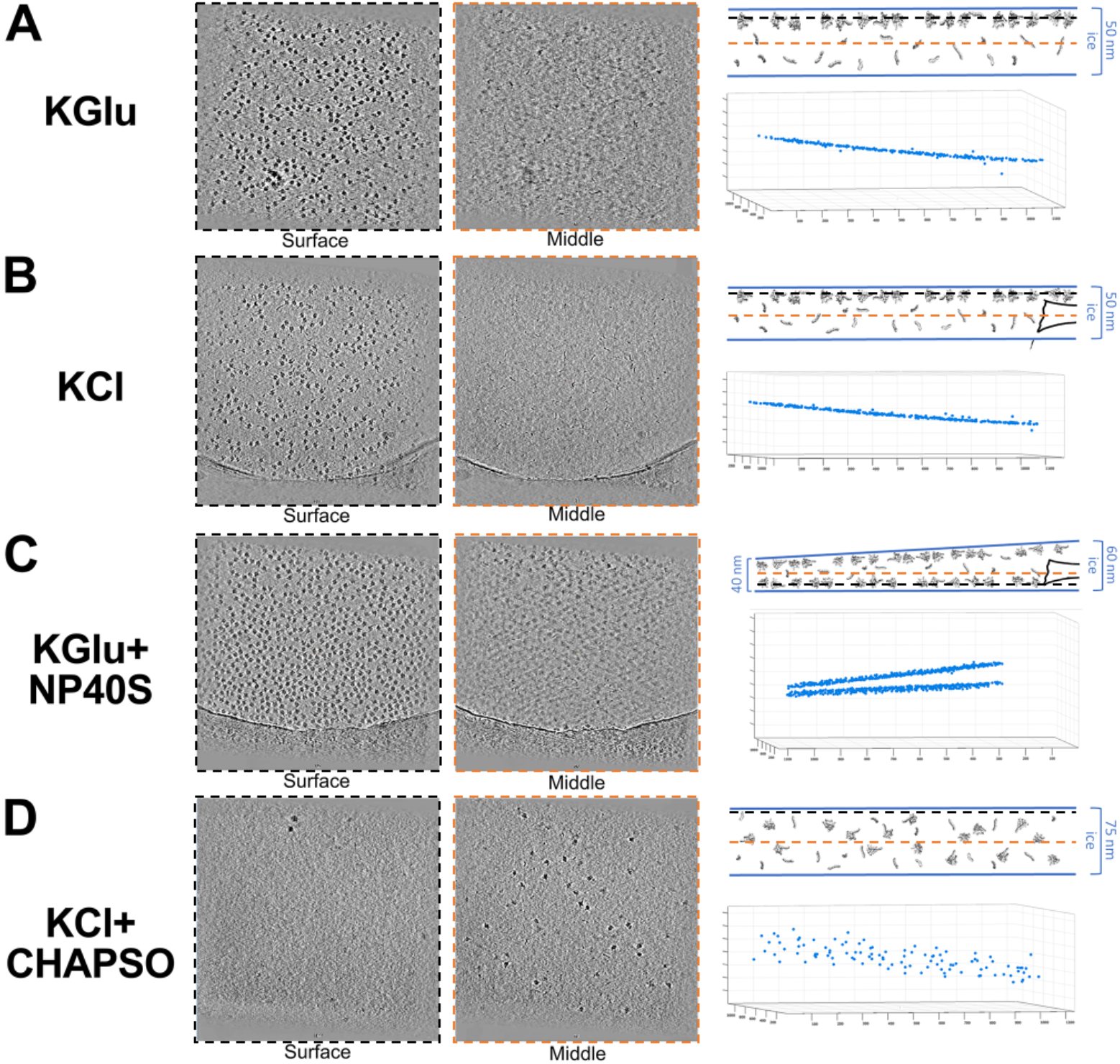
Cryo-ET reveals mechanism for preferred orientation. A – D. (Left Panel) Surface tomographic cross-section of the vitreous ice layer. (Middle Panel) Middle tomographic cross-section of the vitreous ice layer. (Top Right Panel) Schematic diagram of particle distribution in vitreous ice. Top and bottom surfaces of the ice are shown with a solid blue line. Thickness of ice is indicated on the bracket right of the cartoon. 6S-Eσ^70^ and free 6S RNA particles are shown as grey volumes in the cartoon. (Bottom Right Panel) Spatial plot of particles in vitreous ice layer, oriented orthogonal to the ice surface. Each 6S-Eσ^70^ particle is represented as a blue point and graphed based on 3D position in the ice layer. A. Tomogram of 6S-Eσ^70^ particles in KGlu (see Supplementary Table 1). B. Tomogram of 6S-Eσ^70^ particles in KCl. C. Tomogram of 6S-Eσ^70^ particles in KGlu+NP40S. D. Tomogram of 6S-Eσ^70^ particles in KCl+CHAPSO.

The particle orientation distributions and their effects on resulting reconstructions were also analysed using cryoEF [5]. A cross-section through the middle of the expected point spread functions (PSFs) calculated from the particle orientation distributions reveals the severe anisotropy of the KGlu, KCl, and KGlu+NP40S PSFs while the KCl+CHAPSO PSF appeared as roughly a spherical ball (Fig. 1F). Also tabulated in Fig. 1F is the orientation efficiency (*E*_od_) and the fraction of unsampled Fourier space (*f*_empty_) [5], illustrating the dramatic improvement through the use of CHAPSO.

### Cryo-electron tomography shows that orientation bias corresponds to adsorption to an ice surface

We employed fiducial-less cryo-electron tomography (cryo-ET) on the single-particle specimens in order to visualize the locations of the particles in the vitreous ice layer [9,10]. Tilt series of cryo-grids of the 6S-Eσ^70^ complex were collected for each solution condition (Fig. 2). The 6S-Eσ^70^ particles prepared in KGlu and KCl were restricted to a thin layer at one of the ice surfaces (Figs. 2A, B). The complexes were prepared with excess 6S RNA and free 6S RNA molecules could be visualized distributed throughout the vitreous ice layer (Figs. 2A, B, Supplementary videos). The 6S-Eσ^70^ particles prepared in KGlu+NP40S were restricted to two thin layers corresponding to both ice surfaces (Fig. 2C). By contrast, particles prepared in KCl+CHAPSO were excluded from the air/water interfaces and were evenly distributed throughout the middle of the ice layer (Fig. 2D).

### Effect of CHAPSO on particle orientations is concentration dependent

Our results thus far indicate that adsorption of the 6S-Eσ^70^ particles at air/water interfaces gives rise to the severe orientation bias seen in the KGlu, KCl, and KGlu+NP40S samples (Figs. 1A-C, 2A-C). The addition of CHAPSO at the critical micelle concentration (CMC, 8 mM) completely eliminates surface interactions and significantly randomizes the particle orientations (Figs. 1D, 1E, 2D). To investigate if CHAPSO at CMC was required for the full effect, we compared particle orientations for datasets collected with 0, 4 mM (0.5XCMC), and 8 mM (1XCMC) CHAPSO of a different bacterial transcription complex, an *Eco* RNAP ternary elongation complex (TEC) [12]. In the absence of CHAPSO, the particles again exhibited severe orientation bias (Fig. 3A). The particle orientations were significantly spread by the presence of 4 mM CHAPSO (Fig. 3B), but there is a clear difference in the particle spread between 4 mM and 8 mM CHAPSO (Figs. 3B, C) and 8 mM CHAPSO induces a distribution of particle orientations much closer to a random distribution (Figs. 3D, E).

**Fig 3.**
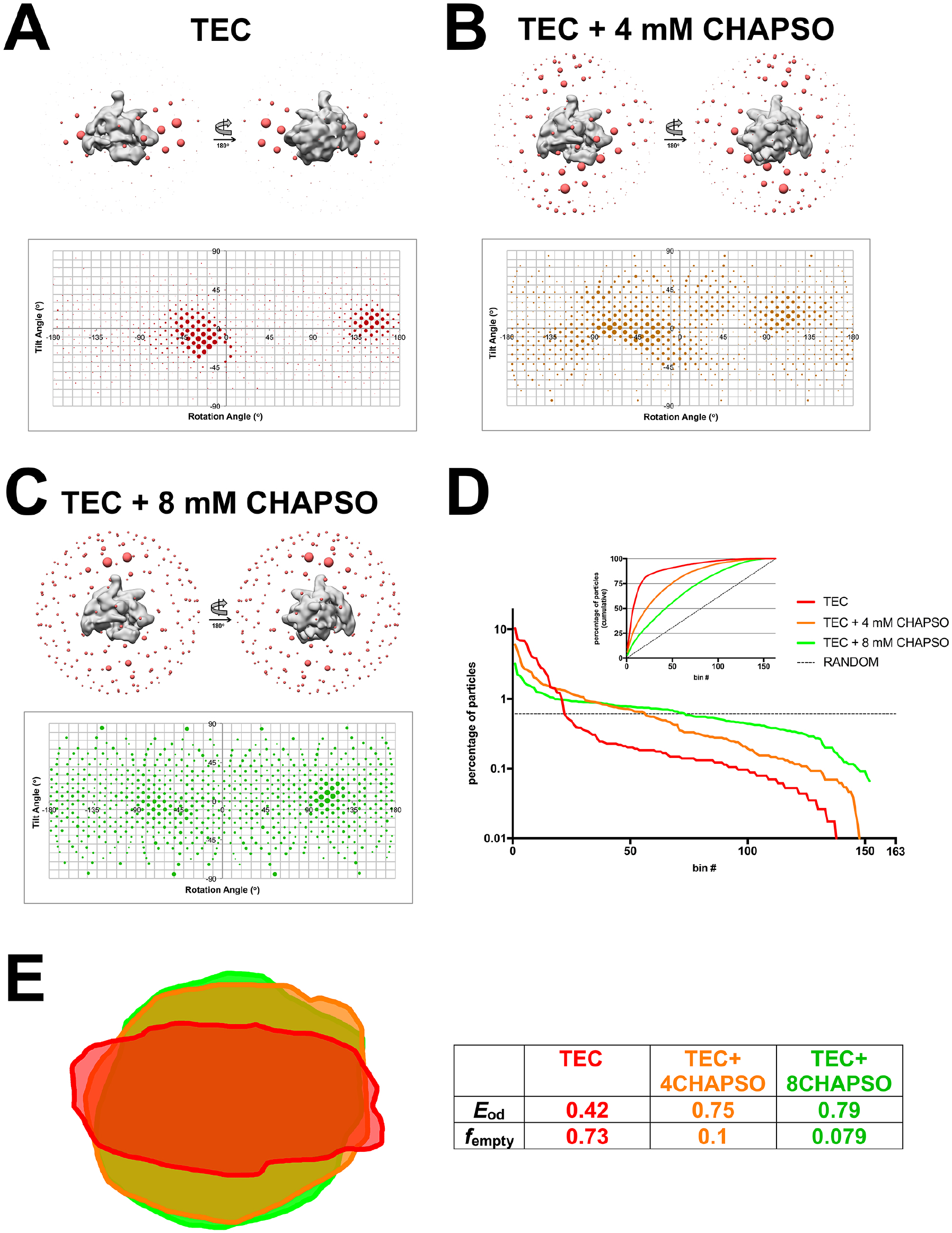
Effect of CHAPSO on particle orientations is concentration dependent. A – C. (Top Panel) 3D distribution plot of particle orientations. Particles were 3D classified into one class using *Eco* core RNAP (PDB ID 4LJZ [14], σ^70^ was deleted and the structure was low-pass filtered using EMAN2) [35] as a 3D template in RELION-2.1 [34]. The resulting density is shown as a solid grey volume and the angular distribution from this alignment is shown as red spheres. Each sphere represents a particular Euler angle and the sphere volume represents the absolute number of particles at that particular angle. (Bottom Panel) 2D distribution plot of particle orientations. Particles are plotted on a tilt angle vs rotation angle graph. Areas of the points represents the percentage of particles at that particular orientation. A. Angular distribution of TEC particles in TEC buffer (20 mM Tris-HCl, pH 8.0, 150 mM KCl, 5 mM MgCl_2_, 5 mM DTT) without CHAPSO. B. Angular distribution of TEC particles in TEC buffer + 4 mM CHAPSO (0.5XCMC). C. Angular distribution of TEC particles in TEC buffer + 8 mM CHAPSO (1XCMC). D. Particles for each dataset were grouped into Euler angle bins (20° rotation angle x 20° tilt angle bind) and then the bins were ranked according to the number of particles populating that bin (bin #1 has the most particles, so on). Plotted on a semi-log scale is the percent of total particles in each dataset by bin #. The horizontal dashed line represents a totally random particle orientation distribution (equal number of particles in each bin). (Inset) Plotted is the cumulative percent particles by bin #. The random distribution is denoted by the dashed line. E. Cross-sections through the middle of the expected PSFs (calculated using cryoEF [5]) are superimposed. Parameters further characterizing the orientation distributions (the orientation efficiency, *E*_od_, and the fraction of unsampled Fourier space, *f*_empty_ [5]) are also tabulated.

### Cryo-EM maps reveal CHAPSO interacts with specific sites on the *Eco* RNAP surface

Examination of the nominal 3.5 Å resolution cryo-EM map of an *Eco* RNAP transcription elongation complex bound to RfaH [13], the highest resolution cryo-EM map available for an *Eco* RNAP transcription complex, revealed three CHAPSO molecules bound to the RNAP surface (Fig. 4A). Retrospective analysis of previously published cryo-EM structures of *Eco* RNAP transcription complexes where 8 mM CHAPSO was used to prevent orientation bias revealed CHAPSO molecules consistently bound at the same sites (Fig. 4B). In each case, the cholic acid-derived amphifacial moiety of CHAPSO was bound to the RNAP while the attached flexible chain and hydrophilic head group were disordered.

**Fig. 4.**
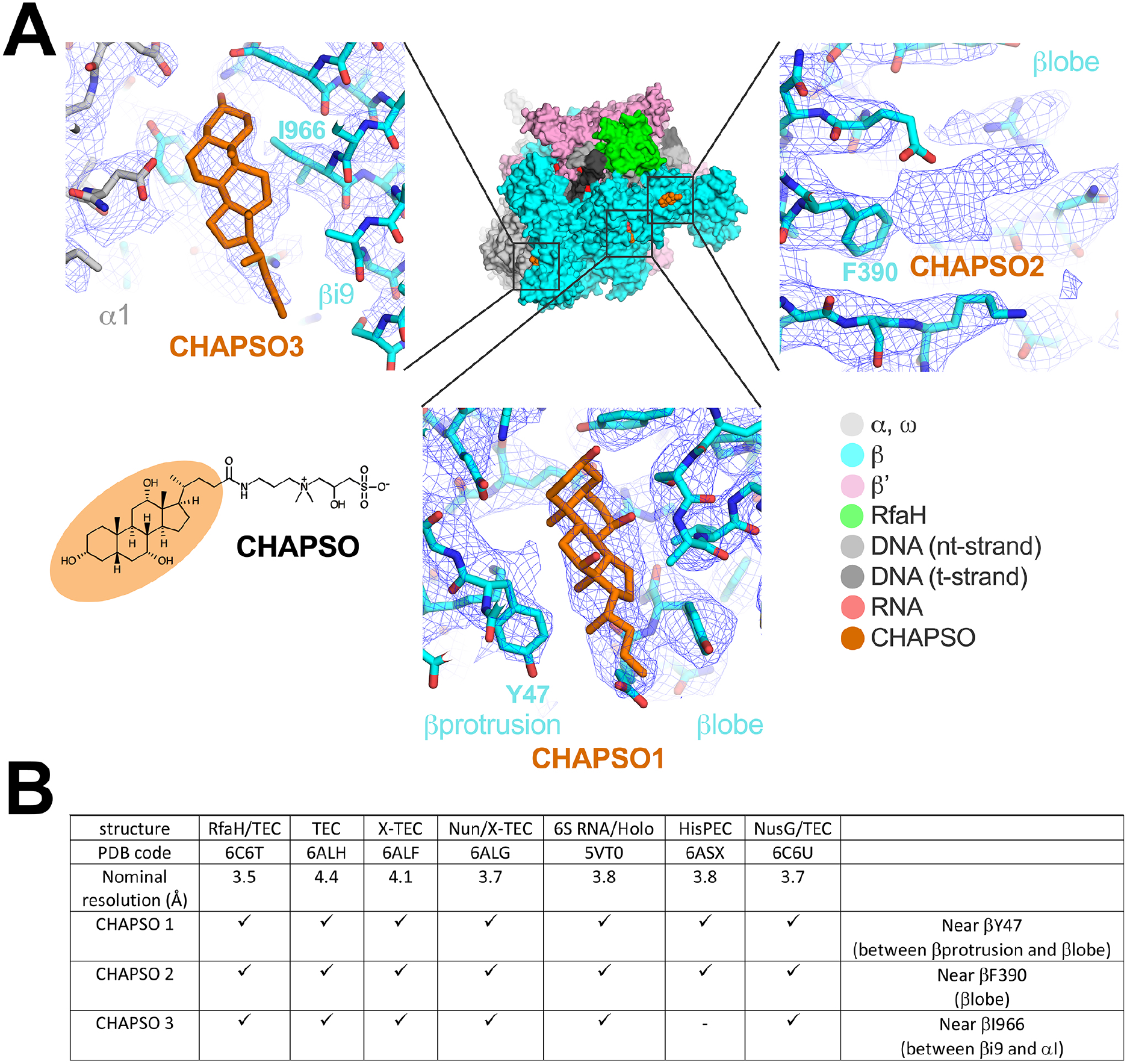
CHAPSO molecules interact with RNAP particles. A. CHAPSO molecules bound to the *Eco* RNAP surface. (top middle) Overall view of the *Eco* RNAP *ops*-ternary elongation complex bound to RfaH (6C6T) [13]. The structure is shown as molecular surfaces color-coded as shown in the color key at the lower right. Shown in orange are three CHAPSO molecules bound to the RNAP surface. (lower left) Molecular structure of CHAPSO. The portion highlighted in orange is resolved in the cryo-EM maps. (top left) Magnified view showing CHAPSO3 along with the nominal 3.5 Å resolution cryo-EM map (blue mesh). (top right) Magnified view showing CHAPSO2 along with the cryo-EM map. The cryo-EM density for CHAPSO2 was not of sufficient quality to determine the CHAPSO orientation. (bottom middle) Magnified view showing CHAPSO1 along with the cryo-EM map. B. Cryo-EM maps of previously published *Eco* RNAP transcription complexes were retrospectively examined for the presence of bound CHAPSO in the three sites. The presence of CHAPSO density in the map is indicated by ‘X’. In the HisPEC (6ASX) [14], a conformational change shifts the position of βi9, disrupting CHAPSO site 3.

## Discussion

From the results of this study, we conclude that:

1. Transcription complexes containing *Eco* RNAP adsorb strongly to air/water interfaces during cryo-EM grid preparation, even in the presence of non-ionic detergents such as NP40S (Figs. 2A-C).
2. The complexes adsorbed to the air/water interface are oriented, confounding single particle reconstruction approaches (Figs. 1, 3).
3. The addition of CHAPSO at the CMC (8 mM) completely prevents adsorption of the complexes to air/water interfaces (Figs. 2D) and dramatically broadens the distribution of particle orientations (Figs. 1D, 1E, 2D, 3C), allowing for the determination of isotropically uniform maps (Figs. 1F, 3E) [5].

Although CHAPSO at 0.5 CMC (4 mM) significantly broadens the particle orientation distribution compared to no CHAPSO (Figs. 3A, B, D) and the calculated PSF and orientational parameters indicate that a detailed reconstruction would be achieved at this condition (Fig. 3E) [5], the full effect on the relative randomization of the particle orientation distribution requires CHAPSO at its CMC (8 mM; Fig. 3). At the same time, due to the high CHAPSO concentration, we observed CHAPSO molecules consistently bound to three sites on the surface of *Eco* RNAP (Fig. 4). Our cryoET results showed that in the presence of 8 mM CHAPSO, the RNAP complexes are completely excluded from the air/water interfaces and found in the middle of the ice layer (Fig. 2D) where rotational diffusion allows for nearly randomized particle orientations. This effect of CHAPSO on eliminating RNAP adsorption at the air/water interface could be due to: 1) CHAPSO at or above the CMC coating the air/water interfaces with a monolayer, altering the interfacial surface properties to prevent RNAP adsorption, and/or 2) CHAPSO molecules binding to the surface of RNAP (Fig. 4), altering the RNAP surface properties to prevent adsorption at the air/water interface. Our result that 8 mM CHAPSO is required for the full effect (Fig. 3) strongly supports hypothesis (1) as the primary factor, since at 4 mM CHAPSO we still observe CHAPSO molecules bound to the RNAP (Supplemental Fig. 2).

In the single particle analyses under conditions where the RNAP complexes are adsorbed to air/water interfaces and oriented, we sometimes observe an orientation distribution comprising one peak (Figs. 1A), while in other cases we observe two peaks, one corresponding to the mirror image projection of the other (Fig. 1B, C). The cryoET analysis revealed some samples where the complexes were adsorbed to only one air/water interface (Figs. 2A, B), while in other samples particles were adsorbed to both the top and bottom interfaces (Fig. 2C), explaining the observation of mirror image projection orientations. While we have not studied this phenomenon specifically, our data suggest that whether the complexes are adsorbed to one or both interfaces is a random occurrence dependent on grid preparation rather than properties of the solution conditions.

A number of cryo-EM structures of bacterial RNAP transcription complexes have benefitted from CHAPSO [7,12-15]. The observation of specifically-bound CHAPSO molecules on the *Eco* RNAP surface (Fig. 4) raises the potential for the introduction of structural artifacts. For this reason it is important to examine the complexes under investigation with sensitive and quantitative functional assays to show that the biochemical function is not altered by the presence of CHAPSO, as was shown in each of the cases listed in Fig. 4B. We believe the very high concentration of CHAPSO (8 mM) allows binding at the observed sites on the RNAP (Fig. 4) but that the binding energy for these sites is very low and is insufficient to alter the conformational/functional properties of the RNAP. For instance, a conformational change in the HisPEC altered the relationship between βi9 and the rest of the RNAP, disrupting CHAPSO site 3 (Fig. 4), and no complexes were observed with CHAPSO bound at this site. In biochemical assays, 8 mM CHAPSO had a minimal (less than 2-fold) effect on the pause lifetime [14].

Orientation bias has long been an issue confounding single particle cryo-EM structure determination [2,3]. Many of the recent advances in high-resolution structure determination of macromolecular complexes by cryo-EM do not address this potential obstacle [5]. A common solution to mitigate particle orientation bias has been to add surfactants during grid preparation [8]. It has often been presumed that particle orientation bias arises from adsorption to the air/water interface and that surfactant additives mitigate the orientation bias by reducing the propensity for interface adsorption. Our cryoET analysis establishes that this is indeed the case for samples comprising *Eco* RNAP transcription complexes and the surfactant CHAPSO (Fig. 2).

Recent studies have established that severe particle orientation bias can be overcome by cryo-EM data collection from tilted grids [6] or by rapid grid freezing after sample application [16]. These are important advances that are potentially generally applicable. Nevertheless, imaging on tilted grids presents many technical obstacles to optimal high-resolution data collection. Moreover, tilting the grid does not address the issue of particle adsorption, which has recently been suggested to cause denaturation for most particles [17]. The specialized plunge-freezing devices necessary for sufficiently rapid plunge-freezing to ‘outrun’ some of the air/water interface adsorption effects are not yet widely available [16]. We show here that the addition of CHAPSO during cryo-EM grid preparation of samples comprising *Eco* RNAP transcription complexes is relatively functionally inert, completely eliminates interaction and orientation of the particles at air/water interfaces (Fig. 2), and significantly broadens the particle orientation distributions to allow determination of high-resolution cryo-EM maps with isotropic PSFs (Figs. 1, 3). These properties greatly facilitate high-resolution structure determination of these complexes using modern cryo-EM approaches [7,12-15].

## Materials and Methods

### Protein expression and purification

*Eco* RNAP TEC was assembled using *Eco* RNAP core lacking the α C-Terminal Domain (ΔαCTD) and was prepared as previously described [12,18]. The 6S-Eσ^70^ complex was prepared as described previously [7].

### Preparation of 6S-/Eσ^70^ for single particle Cryo-EM

Frozen aliquots of *Eco* RNAP ΔαCTD-core and σ^70^Δ1.1 were mixed in a 1:2 molar ratio and incubated for 15 mins at 37^°^C. 6S RNA was added in a 1:1.5 molar ratio and incubated for 15 mins at room temperature. The samples were diluted and detergents added (if used) immediately before grid preparation. 3.5 μL of sample were deposited on glow discharged Quantifoil R 1.2/1.3, 400 mesh, copper grids (EMS) and plunged frozen into liquid ethane using a Cryoplunge 3 system (Gatan).

### Single particle cryo-EM of 6S-Eσ^70^ complex

Cryo-EM grids of 6S-Eσ^70^ were imaged using a 300 kV Tecnai G2 Polara (FEI) equipped with a K2 Summit direct electron detector (Gatan). Dose-fractionated images were collected using UCSFImage4 [19] in super-resolution mode with a nominal magnification of 31,000X, corresponding to a calibrated pixel size of 1.23 Å on the specimen level (0.615 Å for super-resolution). The dose rate on the camera was 8 counts/physical pixel/second using Digital Micrograph (Gatan). The exposure time per movie was 6 s (30 frames) leading to a total dose of 31.7 electrons/Å^2^. Movies were collected using a defocus range from 1 μm to 2.5 μm. Movies were 2X2 binned using IMOD [20] and then drift corrected using MotionCor [21].

### Preparation of 6S-Eσ^70^ for Cryo-ET

For KGlu and KGlu+NP40S conditions (Supplementary Table 1), Eσ^70^ was purified in KGlu buffer using a Superose6 INCREASE column (GE Healthcare). For KCl and KCl+CHAPSO, Eσ^70^ was purified in KCl buffer. Peak fractions corresponding to Eσ^70^ were pooled and concentrated by centrifugal filtration (VivaScience) to 4 mg/mL protein concentration. 6S RNA was added in 1.2 molar excess compared to holoenzyme and incubated at room temperature. Immediately prior to grid freezing, samples of KGlu or KCl were diluted 1:10 with their respective buffers while NP40S was added to the KGlu+NP40S to CMC (0.06 mM) and CHAPSO was added to the KCl+CHAPSO sample to CMC (8 mM). After centrifugation to remove aggregates, 3.5 μL of sample were deposited on glow discharged Quantifoil R 1.2/1.3, Au, 400 mesh grids (EMS) and plunged frozen into liquid ethane using a Vitrobot Mark IV (FEI).

### Acquisition of cryo-electron tomograms of 6S-Eσ^70^

Tilt-series were collected at NYSBC using Titan Krios #1 (FEI Company, Hillsboro, OR) with a Gatan K2 (Gatan, Inc., Pleasanton, CA). Tilt-series were collected using Leginon [22] with 100 ms frames for each tilt image, which were full-frame aligned using MotionCor2 [23]. Tilt-series were collected bi-directionally with a tilt range of −45° to 45° and a tilt increment of 3°. Most tilt-series were collected at a nominal defocus between 4 to 6 microns. Tilt-series were collected with a dose rate around 8 e-/pixel/s and an incident dose of 3.29 e-/Å^2^ for the zero-degree tilt image, with increasing dose for higher tilt angles according to the cosine of the tilt angle, resulting in a total dose of 120 e-/ Å^2^. Most tilt-series were collected at a pixel size of 1.33 Å.

### Cryo-ET data analysis

Tilt-series were aligned using Appion-Protomo [10,24,25]. Tilt-series were first dose compensated using equation 3 in Grant, Grigorieff [26], coarsely aligned, manually fixed if necessary, refined using a set of alignment thicknesses, then the best aligned iteration was reconstructed for visual analysis using Tomo3D SIRT [27,28]. CTF correction was not performed. Subtomogram analysis was performed with Dynamo [29,30]. First, about 10 representative particles were manually picked with two defined Euler angles, averaged together to make a template, then the tomograms were template picked, and the picks were cleaned manually.

### Preparation of *Eco* RNAP TEC for single particle cryo-EM

Purified RNAP ΔαCTD-core was buffer-exchanged over the Superose 6 INCREASE (GE Healthcare Life Sciences) column into 20 mM Tris-HCl, pH 8.0, 150 mM KCl, 5 mM MgCl_2_, 5 mM DTT. At a molar ratio of 1.3:1, template DNA:RNA hybrid was mixed into the eluted RNAP core and incubated for 15 min at room temperature. Subsequently non-template DNA was added and incubated for an additional 10 min [12]. The complex was concentrated by centrifugal filtration (VivaScience) to 3 mg/ml RNAP concentration before grid preparation. CHAPSO was added to the samples to give a final concentration of 0xCMC, 0.5xCMC, or 1xCMC. C-flat CF-1.2/1.3 400 mesh copper grids (EMS) were glow-charged for 15 s. 3.5 μl of sample (~2.0–3.0 mg/ml protein concentration) was absorbed onto the grid, blotted, and plunge-frozen into liquid ethane using a Vitrobot Mark IV (FEI).

### Single particle Cryo-EM of RNAP TEC

Grids of RNAP TEC were imaged using a 300 keV Krios (FEI) (for 0xCMC CHAPSO and 1xCMC CHAPSO datasets) or a 200 keV Talos Arctica (FEI) (for 0.5xCMC CHAPSO). Both microscopes were equipped with a K2 Summit direct electron detector (Gatan). Imaging parameters were as outlined previously [12]. Dose-fractionated images were recorded with Serial-EM [31] in super-resolution mode with a super-resolution pixel size of 0.65 Å (nominal magnification 22,500x and a calibrated pixel size of 1.3 Å) on the Titan Krios and with a super-resolution pixel size of 0.75 Å (nominal magnification 28,000x and a calibrated pixel size of 1.5 Å) on the Talos Arctica. The dose rate at the camera level was 10 electrons/physical pixel/second in Digital Micrograph (Gatan). Images were recorded in dose-fractionation mode with subframes of 0.3 s over a total exposure of 15 s (50 frames). Images were collected over a defocus range of 0.8 μm to 2.6 μm. Movies were gain-normalized and 2X2 binned using ′mag_distortion_estimate′ [32]. Images were drift-corrected and summed using Unblur [26].

### Single particle cryo-EM data analysis

The Cryo-EM data analysis pipeline used for all single particle cryo-EM datasets is illustrated in Supplementary Fig. 1. CTF estimations were calculated for each dataset using Gctf [33]. Particles were picked using Gautomatch (developed by K. Zhang, MRC Laboratory of Molecular Biology, Cambridge, UK, http://www.mrclmb.cam.ac.uk/kzhang/Gautomatch) without a 2D template. Picked particles were extracted from the dose-weighted images in RELION-2.1 [34]. Particles were curated by 2D classification (N classes = 50) and 3D classification (N classes = 3) using a crystal structure of *Eco* RNAP (PDB ID 4LJZ) [11] with σ^70^ removed. The crystal structure was converted to an EM map and low-pass filtered to 60 Å using EMAN2 [35] before classification and refinements. Particles were coarsely aligned to the 3D template using RELION-2.1 [34] 3D classification (N class = 1). Histograms of particle orientations (BILD format) were graphically represented as spheres. The PSFs and orientation distribution parameters (*E*_od_ and *f*_empty_) were calculated using cryoEF [5].

### Accession numbers

Single particle cryo-EM micrographs, cryo-ET tilt-series, cryo-ET tilt-series alignment runs with Appion-Protomo, and cryo-ET tomograms have been deposited to the Electron Microscopy Pilot Image Archive (EMPIAR) with accession codes EMPIARXXXXX.

## Acknowledgements

We thank Z. Li (Harvard Medical School) and T. Walz (The Rockefeller University) for help with EM sample preparation and data collection of 6S-Eσ^70^ and M. Ebrahim and J. Sotiris of The Rockefeller University Evelyn Gruss Lipper Cryo-EM Resource Center for help with data collection. Some of this work was performed at the Simons Electron Microscopy Center and National Resource for Automated Molecular Microscopy located at the New York Structural Biology Center, supported by grants from the Simons Foundation (SF349247), NYSTAR, and the NIH National Institute of General Medical Sciences (GM103310) with additional support from the Agouron Institute (F00316) and NIH (OD019994). A.J.N. was supported by a grant from the NIH National Institute of General Medical Sciences (F32GM128303). This work was supported by NIH R35 GM118130 to S.A.D.

## Supplementary data

Supplementary data to this article can be found online at https://doi.org/X.

**Supplemental Table 1.**
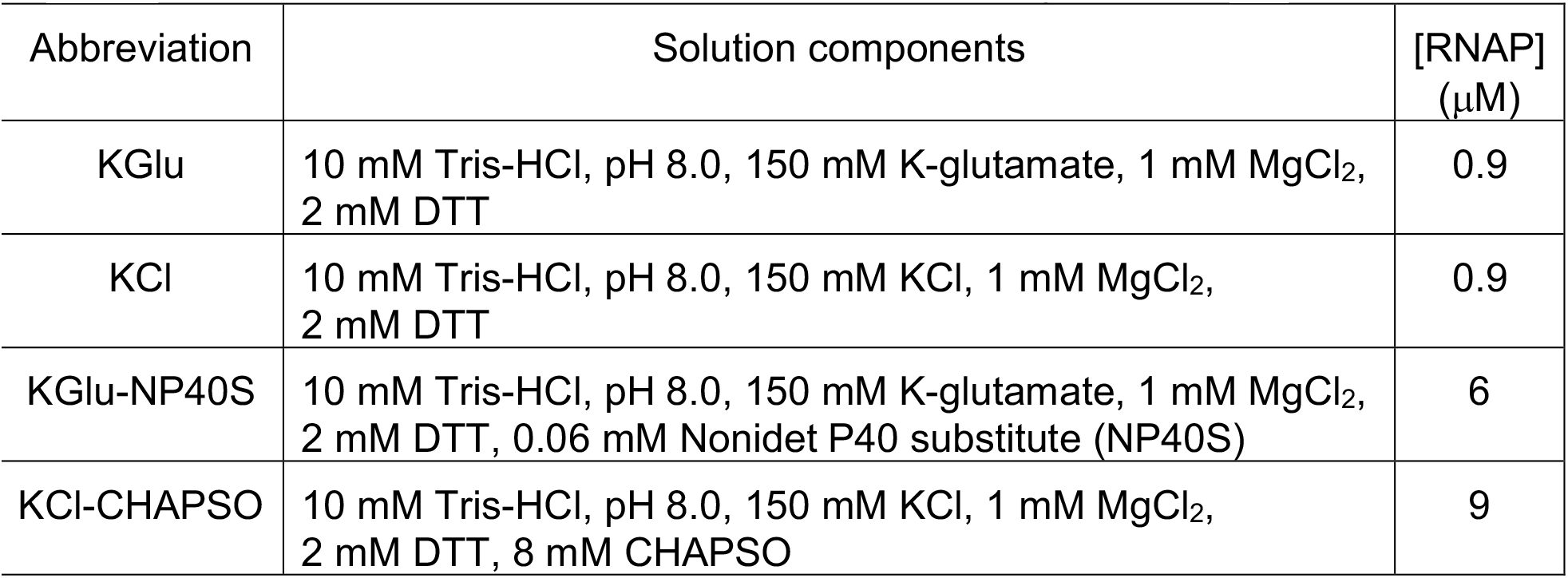
Solution conditions for 6S-Eσ^70^ cryo-EM analyses.

| Abbreviation | Solution components | [RNAP]<br>(μM) |
| --- | --- | --- |
| KGlu | 10 mM Tris-HCl, pH 8.0, 150 mM K-glutamate, 1 mM MgCl<sub>2</sub>,<br>2 mM DTT | 0.9 |
| KCl | 10 mM Tris-HCl, pH 8.0, 150 mM KCl, 1 mM MgCl<sub>2</sub>,<br>2 mM DTT | 0.9 |
| KGlu-NP40S | 10 mM Tris-HCl, pH 8.0, 150 mM K-glutamate, 1 mM MgCl<sub>2</sub>,<br>2 mM DTT, 0.06 mM Nonidet P40 substitute (NP40S) | 6 |
| KCl-CHAPSO | 10 mM Tris-HCl, pH 8.0, 150 mM KCl, 1 mM MgCl<sub>2</sub>,<br>2 mM DTT, 8 mM CHAPSO | 9 |

**Supplemental Fig. 1.**
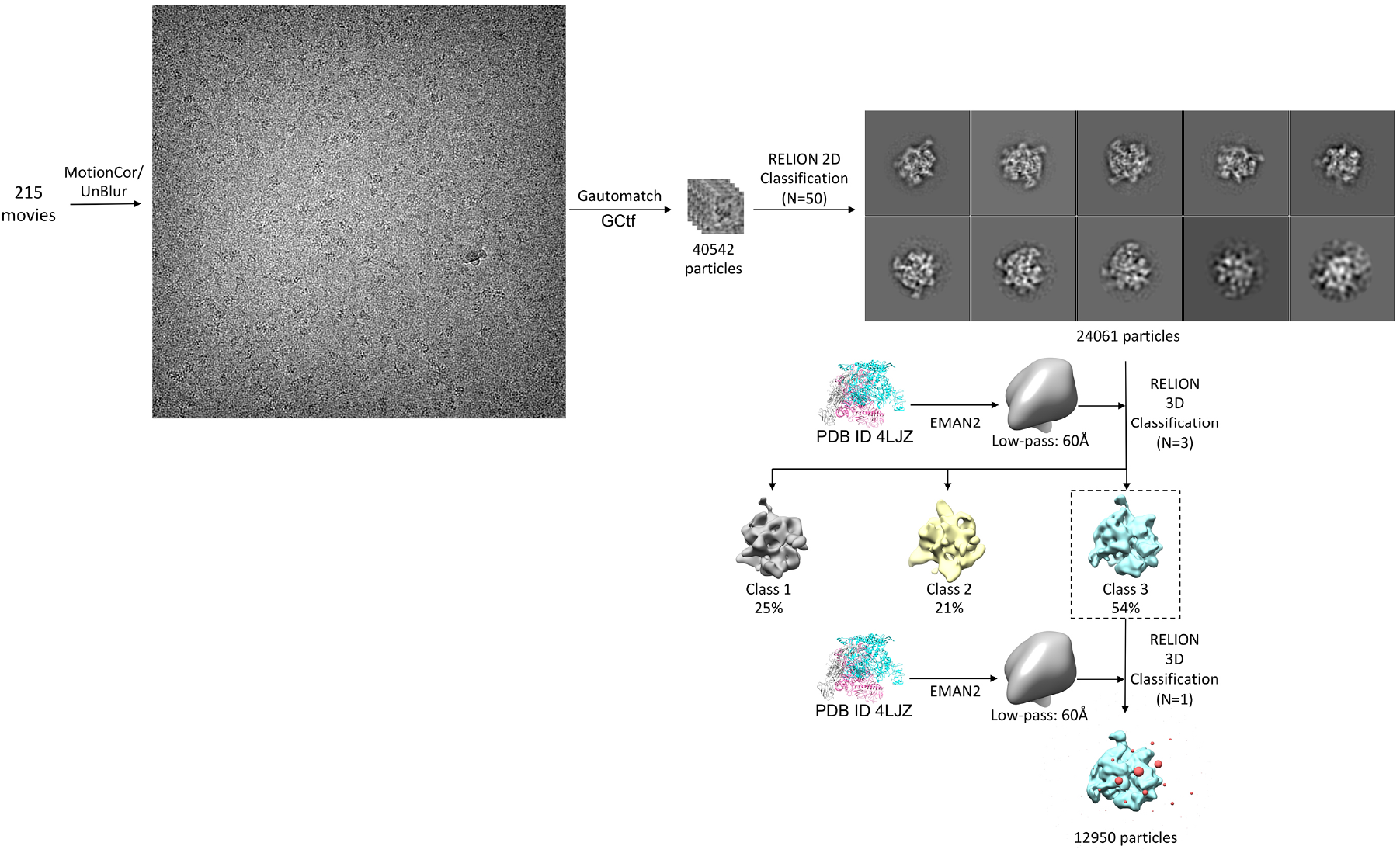
Cryo-EM Data Collection and Analysis of 6S-Eσ^70^ in KGlu. Cryo-EM processing pipeline for 6S-Eσ^70^ in KGlu. This processing scheme was used to process all other datasets in this study. Movies were drift corrected using either MotionCor [21] or Unblur [22] to generate images that were then CTF corrected using GCtf [23]. Particles were autopicked using Gautomatch (K. Zhang, http://www.mrclmb.cam.ac.uk/kzhang/Gautomatch) and cleaned up using RELION-2.1 [24] 2D (N=50) and 3D (N=3) classifications. The final set of particles were aligned using RELION-2.1 [24] 3D classification (N=1) using PDB ID 4LJZ [14] as a 3D template (made in EMAN2 [15]) to determine angular orientations.

**Supplemental Fig. 2.**
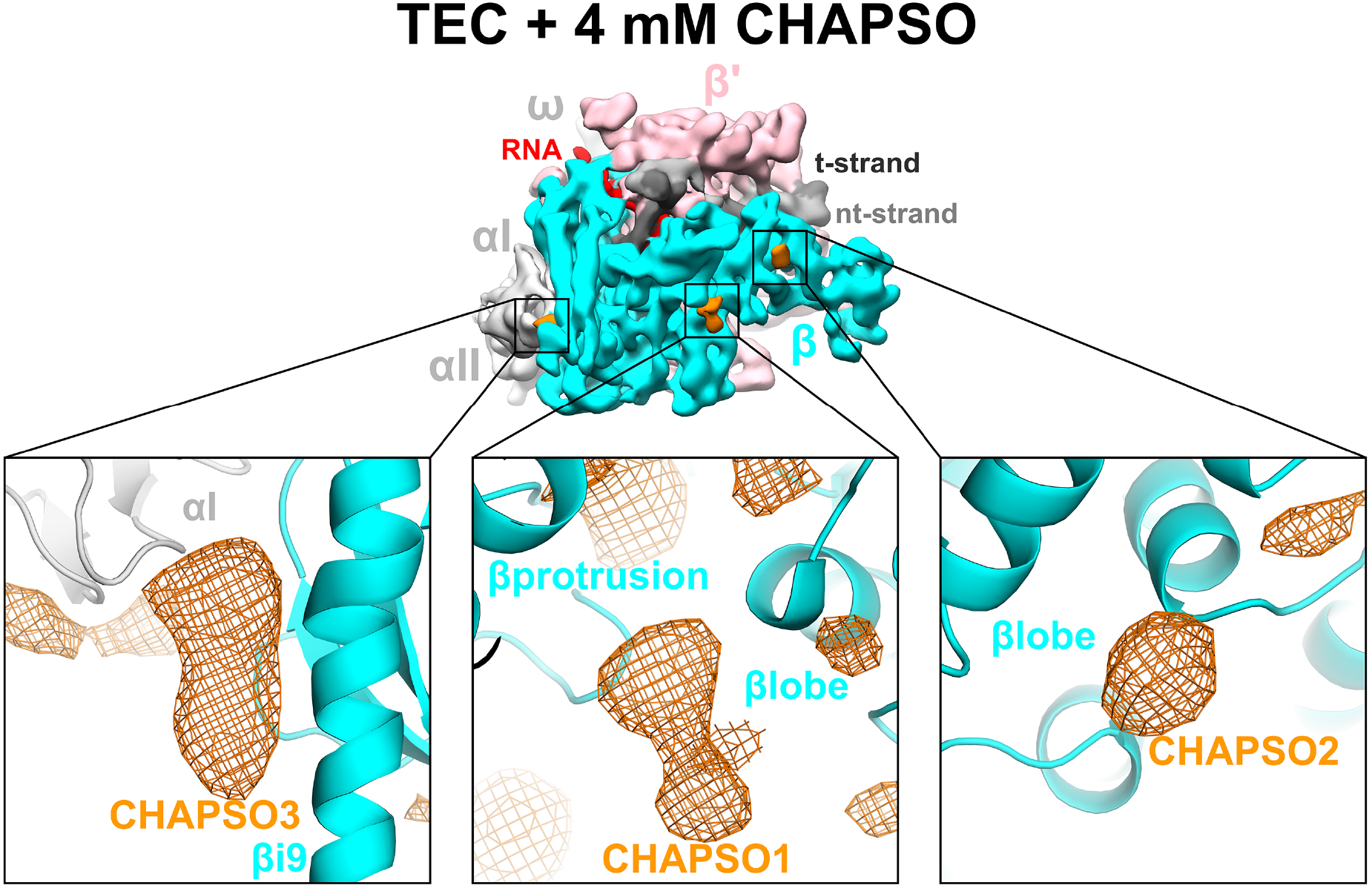
CHAPSO molecules bound to the *Eco* RNAP surface at 4 mM CHAPSO. (top) Overall view of the *Eco* RNAP TEC reconstruction (29,500 particles, nominal 8.3 Å resolution). The structure is shown as molecular surfaces color-coded as follows: αI, αII, ω, light gray; β, cyan; β’, pink; nt-strand DNA, medium gray; t-strand DNA, dark gray; RNA transcript, red. Shown in orange are three CHAPSO molecules bound to the RNAP surface. (bottom) Magnified views of the three CHAPSO sites. The RNAP model is shown as a backbone ribbon. Cryo-EM difference maps (orange mesh) reveal the CHAPSO molecules.

